# Lymphostatin: Structure of a large multi-functional virulence factor

**DOI:** 10.1101/2024.05.21.595110

**Authors:** Matthias Griessmann, Tim Rasmussen, Vanessa J. Flegler, Christian Kraft, Ronja Schneider, Lukas Spantzel, Mark P. Stevens, Michael Börsch, Bettina Böttcher

## Abstract

Enteropathogenic and Enterohaemorrhagic *Escherichia coli* are enteric pathogens of global importance and human infections can be life-threatening. Lymphostatin is a key virulence factor of these bacteria, being required for intestinal colonisation and a potent inhibitor of the mitogen- and antigen-activated proliferation of lymphocytes and proinflammatory responses. In some strains, it also mediates adherence to host cells and influences actin nucleation at sites of attachment. This 365 kDa protein requires glycosyltransferase and cysteine protease motifs for activity against lymphocytes, but high-resolution structural information has proven elusive and the molecular mechanisms by which it acts remain unclear. Here, we describe the structure of lymphostatin from the prototype O127:H6 enteropathogenic *E. coli* strain determined by electron cryo-microscopy. Our results reveal two glycosyltransferase domains, one PaTox-like protease domain, an ADP-ribosyltransferase domain, and a delivery domain. Long linkers act to hold these domains together. These linkers occlude the catalytic sites of the N-terminal glycosyltransferase and protease domains. In this dormant state, lymphostatin binds to HEK-293T cells, where it forms large clusters before being taken up and sequestered into cytosolic foci. With more functional domains than any other known large bacterial toxin, lymphostatin can be regarded as the multifunctional Swiss army knife of pathogenic *Escherichia coli*, enabling complex interactions with the host cells in different environments.

## Introduction

Enteropathogenic and enterohemorrhagic *Escherichia coli* (EPEC and EHEC, respectively) are key causes of pediatric diarrhea and infections can be life-threatening [1]. They belong to a family of attaching and effacing (A/E) bacteria, which adhere intimately to the apical surface of enterocytes and destroy brush border microvilli. EPEC, non-O157 strains of EHEC, and the murine A/E pathogen *Citrobacter rodentium* almost invariably possess lymphostatin (LifA), a 365 kDa protein that plays a pivotal role in intestinal colonisation and has been reported to both inhibit lymphocyte function and mediate bacterial adherence [2,3,4,5,6,7,8]. In affinity purified form, recombinant LifA is active in the femtomolar range against lymphocytes [8,9] and evidence exists that it blocks their proliferation in a manner associated with cell cycle arrest and not apoptosis or necrosis [10]. Lymphostatin homologues exist with similar domains, such as ToxB from *E. coli* O157 which shares lymphocyte inhibitory activity [8].

Lymphostatin exhibits N-terminal homology with the glycosyltransferase domain of large clostridial toxins (LCTs) [3]. It also shares a cysteine protease motif with LCTs that is also found in a wider family of bacterial virulence factors [11]. LCTs play key roles in the pathogenesis of enteric clostridial infections and can drastically alter host cell morphology owing to glycosylation of factors that regulate the actin cytoskeleton [8,9]. Upon binding a surface receptor, LCTs enter the cell via clathrin-mediated endocytosis. Acidification of the endosome induces structural changes and triggers membrane insertion and translocation of the N-terminal portion of the molecule into the cytosol [12]. The cysteine protease domain then mediates autocatalytic cleavage of the protein to release the N-terminal glycosyltransferase (GT) domain into the cytosol, where it subsequently inactivates host Rho GTPases that regulate multiple cellular processes, including actin polymerisation, epithelial barrier integrity, apoptosis and inflammation. The C-terminal part of LCTs is involved in receptor binding and translocation including a delivery domain whose precise role in endosomal escape is not well understood [13]. Uptake and processing of lymphostatin has been hypothesised to occur in a similar way to LCTs [8,14], albeit only the GT domain of LifA belongs to the same protein family as the LCTs, while the delivery domain and the protease domain cluster with other families of protein toxins [15]. Substitution of a DXD motif in the predicted GT domain of LifA abolishes its activity against lymphocytes and binding of uridine diphosphate N-acetylglucosamine, which has been proposed to be the sugar donor molecule [9]. Substitution of the C1480 residue in the catalytic centre of the cysteine protease domain of lymphostatin also abolishes activity, as well as the appearance of an N-terminal 140 kDa fragment in a manner dependent on endosomal acidification [14]. Thus far, only low-resolution structural analysis of lymphostatin has been performed and revealed an L-shaped molecule [9] without resolving the domain architecture.

Here, we have determined the structure of LifA by cryo electron microscopy (cryo-EM), which suggests that LifA forms a catalytically inactive transport form. This form is sufficient to attach to the surface of HEK-293T cells before it is taken up and sequestered into larger cytosolic structures.

## Results

### LifA is a multi-domain protein in two distinct conformations

LifA was stable at pH 4.0 and 6.5 under reducing conditions (Extended Data Figure 1) and allowed for structure determination by cryo-EM. The L-shaped LifA molecule showed an N-terminal arm with abundant α-helices and a C-terminal arm with extended ß-sheets (Figure 1a, Extended Data Figure 2). The two arms were mobile towards one another, with the C-terminal arm adopting one of two distinct conformations. Both conformations co-existed and were not affected by changing the pH from 6.5 to 4.0 (Extended Data Figure 2).

**Figure 1.**
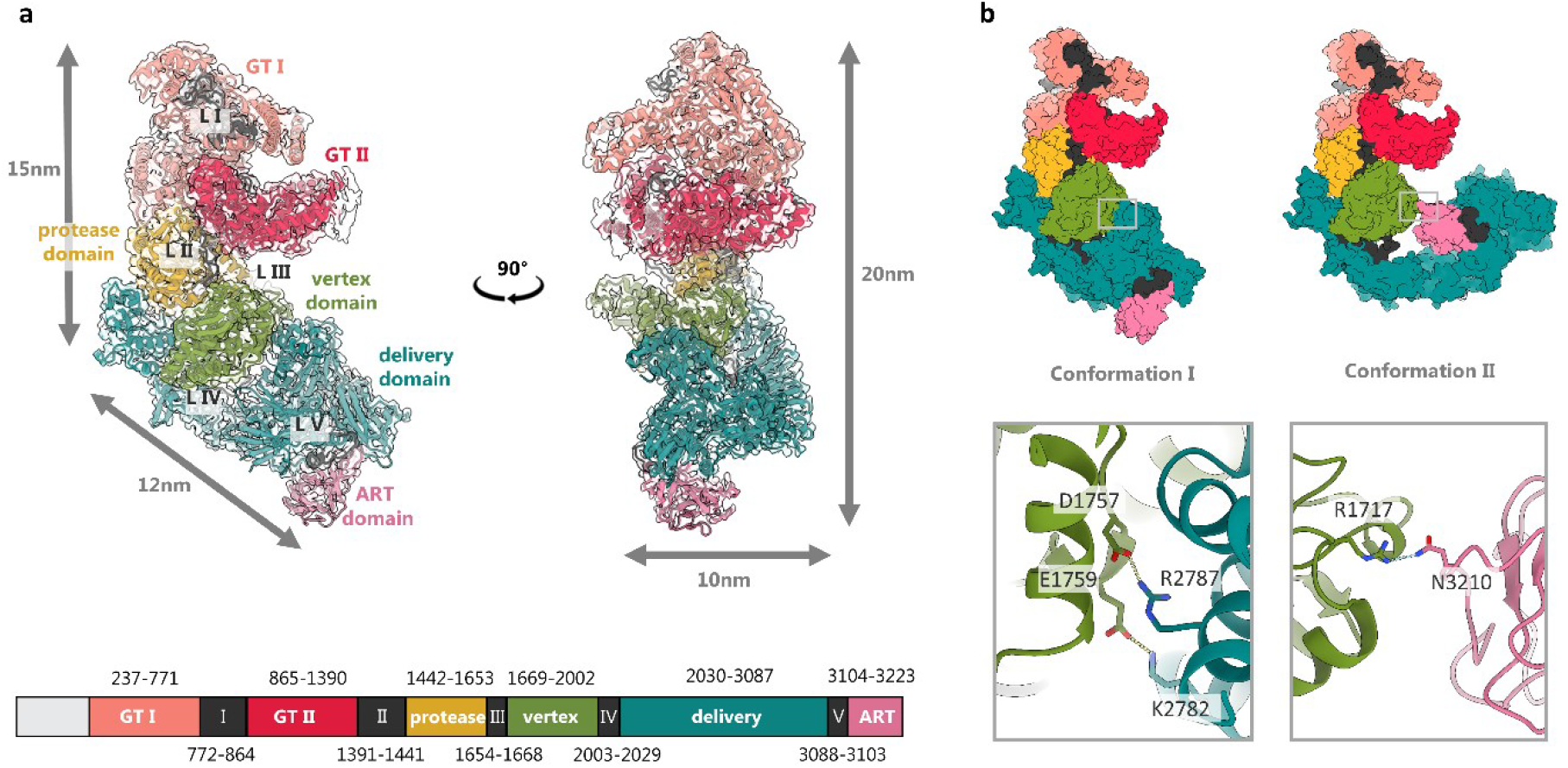
Overall architecture of LifA. **a**, The model of LifA in conformation I fitted into the EM-map of conformation I. The six domains (GT-I, GT-II, protease, vertex, delivery, and ART) are shown in colour and the connecting linkers (LI-LV) in black. The colour code and the residues of the domain boundaries and linkers are indicated in the rectangle below. The map is presented as white, transparent surface. **b**, Surface representation of the models of conformation I and conformation II in the same orientation. The rectangles mark the interaction site between the vertex domain and the delivery domain in conformation I and between the vertex domain and the ART in conformation II. The close-ups below show the respective contact points between the two domains. In conformation I there are two salt bridges (D1757-R2787 and E1759-K2782) and in conformation II one hydrogen bond (R1717-N3210).

We identified six domains in LifA connected by five linkers (L-I to L-V; Figure 1a, Extended Data Figure 3). The N-terminal arm had the GT44 domain (GT-I) (InterPro PF12919) [16] at its tip and the C58_PaToxP-like protease domain (InterPro CD20495) [16] located near the vertex (Figure 1). Another GT44 domain (GT-II) (InterPro PF12919) [16] was found between these two domains. The vertex of LifA was composed of an α-helical domain and had no similarity to other proteins of known structure and function. The C-terminal arm was mainly composed of a region with ß-sheets, which is classified as a domain of unknown function DUF3491 (InterPro PF11996) [16]. This domain is characteristic of the LifA, Efa-1 and ToxB-like protein family [17] and is designated as an evolutionary conserved translocation and delivery apparatus in LCT-like toxins [18]. Therefore, we will refer to it as the delivery domain. Additionally, an ADP-ribosyltransferase (ART) domain was located at the end of the C-terminal arm and folded either beneath the delivery domain in conformation I or connected with the vertex domain after rearrangement of the delivery domain in conformation II (Figure 1b).

### LifA has two glycosyltransferase domains

The GT-I domain is structurally closely related to the clostridial TcdA GT-domain (Extended Figure 4a). The latter is well characterized by both X-ray crystallography and cryo-EM [19,20,21,22,23]. A characteristic DXD motif in TcdA coordinates a Manganese-ion that stabilizes the leaving phosphate group of the sugar substrate. In GT-I of LifA this functionally essential motif [9] was found in a pocket at residues 557-559 (DTD, Figure 2a). On the opposite side of this pocket was an LNG motif (residues 667-669, Figure 2A), which is responsible for substrate binding in TcdA and therefore marked the likely binding site of the sugar moiety of UDP-GlcNAc [9]. The substrate binding site of GT-I was occupied by a loop (residues 532-542, Figure 2a). In addition, the entry to the binding cleft was blocked by the linker (L-I) that connected GT-I with GT-II (residues 772-864). L-I wrapped around the whole GT-I domain and filled the grooves on the surface of the domain (Figure 3b).

**Figure 2.**
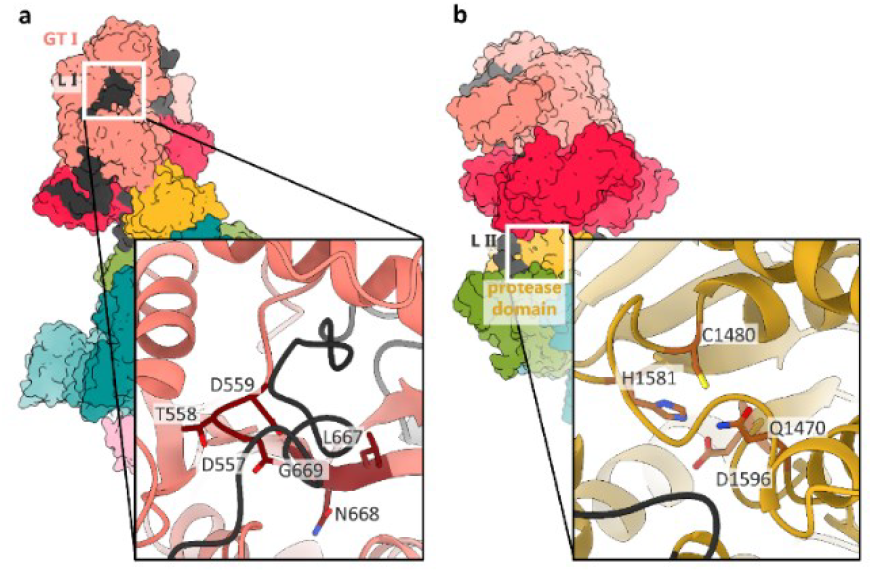
Active sites of GT-I and the protease domain. Surface representation of the model of LifA in conformation I with close-ups of the active sites in cartoon representation. **a**, The close-up of the GT-I shows the characteristic DXD-motif for catalysis (D557, T558 and D559) and the LNG motif (L667, N668, G669) for substrate binding. The entrance to the active site is blocked by L-I (black). **b**, The close-up shows the catalytic triad (C1480, H1581 and D1596) of the protease domain. The loop connecting the functional residues Q1470 and C1480 blocks the entry to the active site. L-II (black) is a possible target for auto-cleavage but requires major structural rearrangements to move into the active site.

**Figure 3.**
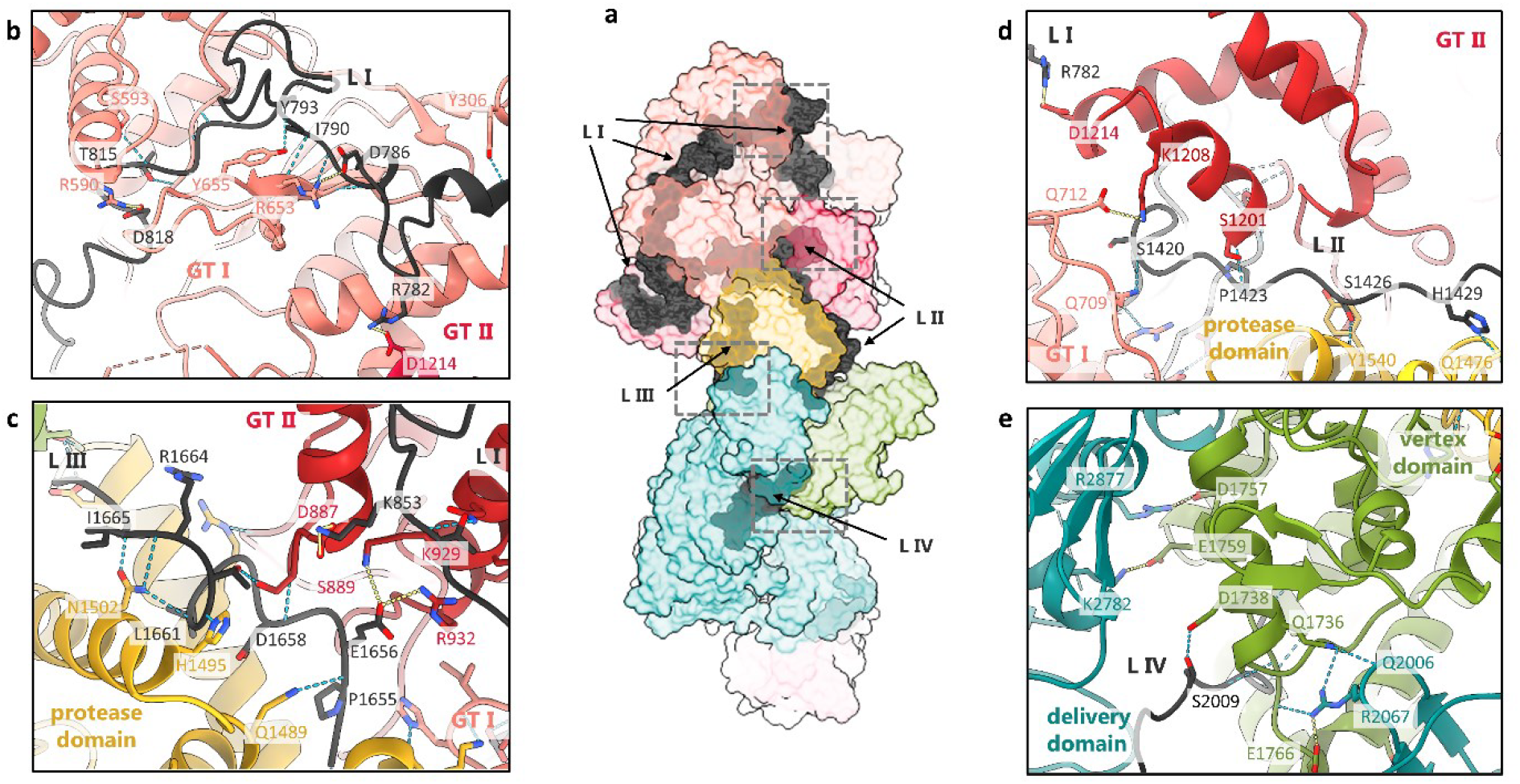
Domain-domain and linker-domain interactions in LifA in conformation I. **a**, Model of LifA in surface representation with close ups of the linker regions (**b-e**) indicated by squares in **a.** The close-ups show the model in cartoon representation. The side chains of hydrogen bonded residues are shown together with the label of the residue number and the hydrogen bond as dotted line. The domains are coloured the same as in Figure 1. The orientations in the close-ups (**b-e**) differ from the overview in **a**. The linkers are at the interfaces between domains and interconnect several domains by hydrogen bonds.

The GT-II domain mapped to residues 865-1390. FoldSeek [24] identified the GT-domain of *Clostridioides difficile* toxin B (TcdB) as the closest homologue with known structure. However, the sequence conservation was low (13% identity), which explains why this GT domain was not previously identified via sequence-based analyses. Notably, the GT-II domain lacked a DXD motif, instead showing an EEN motif in its place (residues 1192-1194, Extended Data Figure 4b). In contrast to the DXD motif in GT-I, the EEN-motif was fully exposed on the surface in a shallow indentation (Extended Data Figure 4b).

### The active site of the LifA protease domain is inaccessible

The strictly conserved catalytic residues C1480, H1581 and D1596 constitute the active site of the LifA C58_PaToxP-like protease domain with Q1470 stabilizing reaction intermediates (Figure 2b) [14]. As in the GT-I domain, the active site was shielded from the surrounding solution by a loop between the functionally relevant residues Q1470 and C1480. The loop was wedged between the active site and one of two long helices that cross each other. These helices were structurally reminiscent of prodomains of papain-like proteases (prodomain-like helices, Extended Data Figure 4c) that regulate the protease activity [25]. The prodomain-like helices were hydrogen-bonded to the GT-II domain, the vertex domain, the functionally relevant loop, and the linkers preceding (L-II) and following the protease domain (L-III; Figure 3c). Thus, the prodomain-like helices were tightly fixed in their position and could not move aside to release the functionally important loop from the entrance to the active site of the protease.

Analogy to the LCTs suggests that the protease domain could release the N-terminal effector domains by autocatalytic cleavage [26]. A potential cleavage target in LifA is the linker that connected the protease domain with the upstream GT-II domain (L-II, residues 1391-1441). L-II was close to the active site of the protease domain but too distant for direct cleavage (Figure 2b). It formed several hydrogen bonds with GT-I, GT-II, the vertex domain, and the prodomain-like helices (Figure 3). As such L-II glued the domains together and held LifA in a compact state. At the same time the hydrogen bonding network limited the ability of the linker to move into the active site of the protease. The cleavage of L-II would release an approximately 166kDa fragment that includes both GT domains and is similar in size to the fragment found in LifA-treated bovine T lymphocytes [14].

### Conformation I and II differ by rearrangements in the delivery domain

The delivery domain was mainly composed of ß-strands and was subdivided into four ß-sandwich sub-domains (S1-S4, Extended Data Figure 5). The N-terminal ß-sandwich (S1) was the largest, containing 42 strands (residues 2060-2629) while S2-S4 had only 7-10 strands. Such ß-sandwiches are typical for bacterial translocation and pore forming systems, as well as receptor binding and adhesion domains [12,27,28,29]. The relative arrangement of S2-S3 changed between conformation I and II. In conformation I, S2 formed two salt bridges with the vertex-domain (E1759-K2782 and D1757-R2877, Figure 1b). This interaction was lost in conformation II, and instead, the ART domain made one hydrogen bond to the vertex domain (R1717-N3210, Figure 1b). The underlying domain rearrangement involved a rigid body rotation around G2630 of the downstream S2-S4 and the ART-domain. This drastic domain rearrangement transformed LifA from a compact form into an elongated form. At pH 4.0 and 6.5 both conformations were present (Extended Data Figure 2). This differs from TcdA and TcdB LCTs, which transition from a compact form into an elongated form when the pH is lowered.

### Fluorescence-labelled LifA clusters on the membranes of HEK-293T cells

To gain insight into the interaction between LifA and living cells, we used HEK-293T cells stably expressing the Neurotensin receptor 1 fused to mNeonGreen (NTSR1-mNG) [30]. These cells were exposed to LifA labelled with the fluorescent dye Cy3B (LifA-Cy3B). NTSR1-mNG serves the purpose of highlighting the plasma membrane of the cells for dual-colour confocal fluorescence lifetime imaging (FLIM) in a microfluidic chamber (Figure 4, Extended Data Figure 6). Approximately 20 min after addition, LifA-Cy3B started to form fluorescent clusters on the cell surface, which slowly grew within the first two hours (Extended Data Figure 6d).

**Figure 4.**
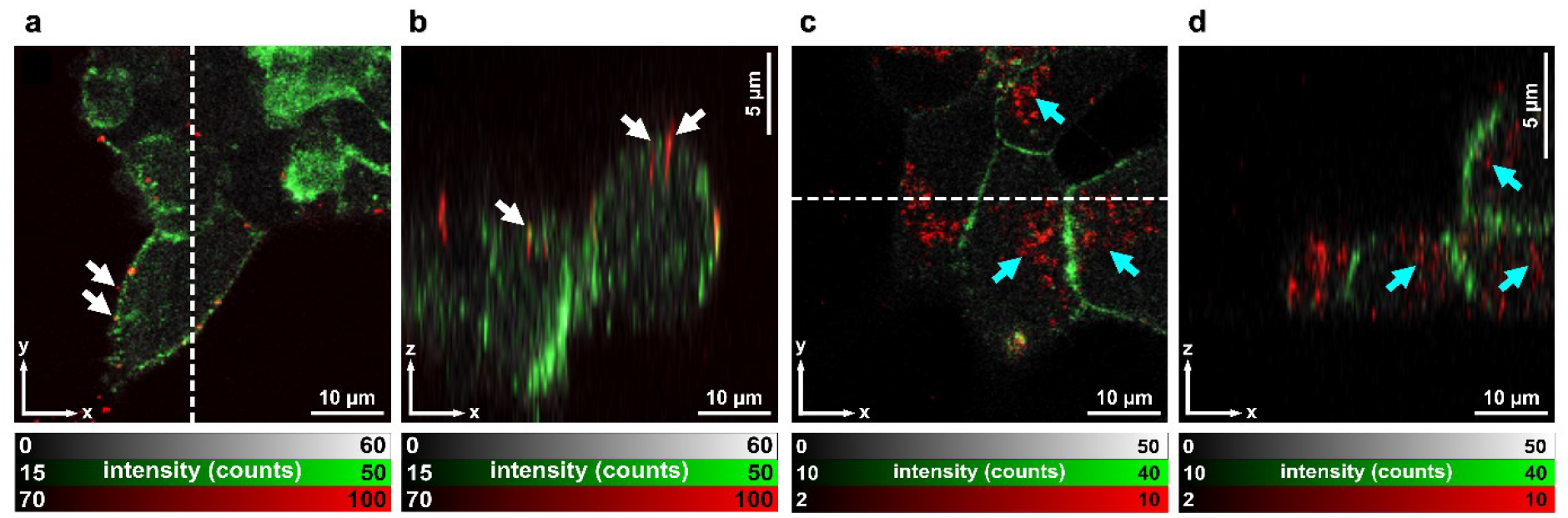
LifA clusters on HEK-293T cell membranes and is internalised in living cells. **a**, LifA formed clusters (white arrows) on the plasma membranes of live HEK-293T cells after incubation with LifA labelled with Cy3B. Confocal z-stack FLIM images were recorded 30 min after LifA-Cy3B addition to the cells. LifA-Cy3B is shown in red, plasma membrane-embedded NTSR1-mNG is shown in green. Fluorescence intensities are shown in grey. **b**, No internalisation of LifA-Cy3B could be observed within 90 minutes. **c**, Incubation for 15 h with LifA-Cy3B showed completed uptake of the protein into living HEK-293T cells (cyan arrows). **d**, LifA sequesters to bright clusters inside the cells.

The clusters adhered tightly to the plasma membranes and could not be removed by extensive washing (Figures 4a, 4b).

To study the localisation of LifA over longer time periods, we incubated HEK-293T cells for [Grab your reader’s attention with a great quote from the document or use this space to emphasize a key point. To place this text box anywhere on the page, just drag it.]

15 h with LifA-Cy3B in cell culture medium. Before measuring FLIM in the microfluidic chamber, cells were washed with PBS buffer. Afterward NTSR1-mNG was localised at the cell surface. In contrast, most of the fluorescence associated with LifA-Cy3B was found in bright foci inside the cells. This suggests that most of the internalisation of the protein was completed at the time (Figures 4c, 4d, Extended Data Figure 6c and d).

## Discussion

Lymphostatin is one of the largest proteins yet described in *E. coli* and possesses multiple structural domains. It has the characteristic topology of an AB exotoxin with a toxic A part consisting of the functional active domains for the cytotoxic effect on the host cells and a structurally more variable B part involved in the host cell entry and endosomal escape. Lymphostatin follows the prototypic domain architecture of an LCT-like toxin with a GT-I domain, a protease domain, and a delivery domain. Though, it also diverges from this by an additional GT-II domain and an ART domain. While the GT-II domain and the GT-I domain are close together in the A-part, the ART domain is at the other side of LifA in the B-part indicating activity of the ART domain in a different spatial and temporal context. ART domains are effectors in many bacterial toxins such as Cholera-like toxins, Diphtheria-like toxins, C2-like binary toxins, C3-like toxins [31]. However, the position in the B-part and the combination with a GT-domain in a single toxin are atypical. Bacterial toxins often adopt inactive transport forms to enable safe delivery to the target without collateral damage at other places. Along these lines, we found that the GT-I domain and the protease domain are in an inactive state, with the entry to the substrate binding site blocked. In both cases, this blockage is due to an intra-domain loop and an extra-domain linker occluding the substrate binding pocket. Therefore, activation necessitates global conformational changes. These changes involve the loss of the extended hydrogen bonding network of the linkers to the surrounding domains, as well as more confined conformational changes within the domain.

In contrast to GT-I, the substrate binding site of the GT-II domain is unprotected although GT-II is most likely inactive due to the absence of a catalytically important DXD-motif. Instead, the functional relevance of GT-II could be the pre-binding of the potential substrates in the proximity of the functional GT-I domain. The binding site in GT-II is enclosed from the opposite side by the mobile, N-terminal helices of GT-I (Extended Data Figure 7). These N-terminal helices of GT-I are a membrane localization domain of the GT-domain in LCTs [32]. Binding larger target proteins to GT-II and/or associating GT-I with the membrane may separate the two domains and release L-I for GT-I activation. This mechanism ensures that GT-I is only activated when target substrates are present at the target site.

Following the assumption that lymphostatin is processed in a way similar to the LCTs TcdA and TcdB, we observed that endosome acidification is required for C1480-dependent cleavage of the protein inside cells [14]. In TcdA and TcdB the C-terminal combined repetitive oligopeptides (CROPs) domain restructures at low pH and releases the delivery domain (IPR024769, TcdA/TcdB_pore_forming) [20,28]. Such CROPs are absent in lymphostatin, and its delivery domain belongs to a different protein family requiring a different mechanism of action. Along these lines, we do not see pH dependent changes in the conformation suggesting divergence from the structural maturation of LCTs. However, we could confirm uptake of lymphostatin by HEK 293T cells into vesicle-like structures (Figure 4c, 4d).

Lymphostatin does not induce apoptosis or necrosis in cells, but it does arrest the host cell cycle [10] and has been implicated in bacterial adherence to host cells [3,5]. It is conceivable that lymphostatin aids focal bacterial adherence as evidenced by the formation of large stable clusters on the cell surface that resist extensive washing. Immunofluorescence microscopy using antibodies against the protein found it to be enriched at the perimeter of bacterial cells in a manner similar to the bacterial adhesin intimin [5]. Multiple clustered LifA molecules offer interconnected interaction sites for multivalent binding with high avidity for both the host cell and the attaching bacteria. The delivery domain (DUF3491) is the likely responsible for adherence to the host cell due to its ß-sandwich domains, which are typical for receptor binding domains. However, the primary receptor on the host cells remains unknown.

It is noteworthy that while affinity purified lymphostatin can inhibit lymphocyte function in isolation, during infection it may also be directly injected into cells. Evidence exists that it can be secreted by a Type 3 secretion system in EPEC, the function of which is to inject bacterial proteins directly into enterocytes and is critical for intestinal colonisation [17]. Consequently, lymphostatin may interact with host cells in distinct ways, with autocatalytic cleavage following endosomal uptake and membrane insertion being one pathway, and direct injection into the cytoplasm being another. At least some of the protein also appears enriched at the bacterial surface and influences attachment [5]. Lymphostatin was the first of a wider family of lymphocyte inhibitory factors to be identified. ToxB from *E. coli* O157 shares its glycosyltransferase, cysteine protease and delivery domains and is similarly able to inhibit lymphocyte proliferation and pro-inflammatory cytokine synthesis [8]. Moreover, sequencing of the genome of the EPEC strain in which lymphostatin was first described (strain E2348/69) has revealed that it contains a truncated LifA-like gene [33] which was required for efficient formation of A/E lesions on human intestinal explants, at least when other Type 3 secreted effectors were absent [34]. Our high-resolution structure of lymphostatin has revealed an unprecedented number of functional domains in a single protein and provides rich information to interpret the potential impact of differences between lymphostatin and its homologues, both within the same strain and between pathotypes and bacterial species. It is noteworthy that *lifA* and *lifA*-like genes are encoded on integrative elements in E2348/69 together with other effectors of the bacterial Type 3 secretion system [33]. Recombination and horizontal gene transfer events are therefore likely to have shaped its remarkable architecture and multi-functionality.

## Experimental Procedures

### Protein Expression and purification

Recombinant LifA containing a C-terminal His-tag in the pRham Vector (*Lucigen*) [9] was overexpressed in E. cloni® 10G cells (*Lucigen*). All cultures contained 50 μg/ml kanamycin for bacterial selection. After overnight cultures (LB medium with 0.5 % (w/v) L-glucose), cells were transferred to fresh LB medium with 0.25 % L-glucose and grown up to A_650nm_ 0.8. The cells were then transferred to glucose-free TB medium containing 2 mM MgCl_2_. At A_650nm_ 2.0, the temperature was reduced to 30 °C for 1 h and then expression was induced with 0.2 % (w/v) L-rhamnose. After 3 h, cells were harvested by centrifugation (3857 g for 27 min) and the pellet was equally distributed to four 50ml tubes before being shock frozen into liquid nitrogen and stored at -80°C until further use.

For purification, one tube of shock-frozen cell pellet (containing approx. 15 g) was re-suspended in 40 ml lysis buffer which contained 20 mM sodium phosphate buffer pH 6.3, 300 mM NaCl, 500 mM 3-(1-Pyridin)-1-Propansulfonat (NDSB-201; *Sigma*) for protein stabilisation, 20 mM imidazole, 10 % (v/v) glycerol, 100 μM phenylmethanesulfonyl fluoride (PMSF), 0.1 % (v/v) Tween-20, 1 mM 2-Mercaptoethanol, one tablet of EDTA-free cOmplete™ Protease Inhibitor Cocktail (*Roche*; *Sigma Aldrich*) per 3 g of cell pellet, 2 mg DNase (*Roche*) and 10 mM MgCl_2_. Cells were lysed by two passages through a benchtop cell disruptor (LM-20 Microfluidizer®; *Microfluidics*) at 1.25 kbar. The lysate was clarified by centrifugation (7197g for 30 min) and applied twice at 4 °C to a Protino Ni-IDA 2000 column (*Macherey-Nagel*), pre-equilibrated with IMAC buffer containing 50 mM sodium phosphate buffer pH 8.0 and 300mM NaCl. LifA was eluted in 1 ml fractions with IMAC elution buffer that contained 250 mM imidazole and was adjusted to pH 6.5. LifA was further purified by size exclusion chromatography using a Superdex 200 10/30 GL increase column (*Cytiva*) pre-equilibrated with SEC buffer (25 mM sodium phosphate pH 6.5, 150 mM NaCl, 75 μM MnCl_2_, 5 % (v/v) glycerol, 1.5 mM TCEP). LifA peak fractions (Extended Data Figure 1c) were pooled and concentrated to approx. 0.9 mg/ml with a 100 kDa cutoff Amicon Ultra centrifugal filter (*Millipore, Merck*) while glycerol was removed via washing with concentration buffer (25 mM sodium phosphate pH 6.5, 100 mM NaCl, 1 mM Uridine diphosphate N-acetylglucosamine (UDP-GlcNAc), 75 μM MnCl_2_, 1.5 mM TCEP). The protein concentrations were determined photometrically at A_280nm_ either with the build-in detector of the HPLC system (NGC™ Chromatography system, *Biorad*) or with a spectrometer (Genesys 50, *Thermo Fisher*). For samples to be vitrified at pH 4.0, the concentration buffer without UDP-GlcNAc was adjusted to pH 4.0 with HCl before use.

### Protein Analysis by SDS-PAGE and Western Blotting

Protein quality was assessed by SDS-PAGE and Western Blot. Samples were mixed with Laemmli Sample Buffer (5 % ß-mercaptoethanol, 0.02 % bromophenol blue, 30 % glycerol, 10 % SDS, 250 mM Tris at pH 6.8) and subsequently incubated at 70°C for 10 minutes before being applied to the gels. The acrylamide gels were either directly stained with Coomassie (*Roth*) or used for Western Blot. Blotting onto nitrocellulose membranes (*Cytiva*) took place at 4 °C in Transfer Buffer (24 mM TRIS, 194 mM Glycine, 0.05 % SDS, pH 8.3) for approx. 16 h with low concentrated acrylamide gels (7,5 %) to ensure complete transfer of large proteins. Membranes were blocked for 1 h with 3 % (w/v) milk powder solved (*Roth*) in TBST buffer (150 mM NaCl, 50 mM Tris at pH 7.6, 0.1 % Tween-20) before being washed with TBST. Then, membranes were incubated under soft shaking with Penta-His antibody (Penta His HRP Conjugate; *Qiagen*) for 1 h with 0.5 % (w/v) milk powder in TBST. The final antibody dilution was 1:50,000. After washing of membranes, pre-mixed Western Blot solution (1:1 mixture of peroxide solution and luminol enhancer solution, both *Thermo Fisher*) was applied. The membranes were imaged with a Fusion FX Imaging chamber (*Vilber*) with detection times of 5 to 10 minutes (Extended Data Figure 1a).

### Fluorescence labelling of LifA with Cy3B

For labelling of LifA with the fluorescent dye Cy3B maleimide (*AAT Bioquest*), washing and elution during IMAC were performed at pH 8.0 to have LifA in suitable buffer for the subsequent conjugation reaction. 400 μl of the main IMAC elution fraction was concentrated (Amicon Ultra, 100 kDa MWCO) to concentrations between 1 and 2 μM to improve labelling efficiency. TCEP concentrations were increased to between 2 and 5 μM before labelling. Labelling took place for 15 minutes at room temperature with 15x molecular excess of dye to protein before further purification via SEC as described above (Extended Data Figure 1d). Separation of unbound dye and the absence of oligomeric LifA-Cy3B was confirmed by fluorescence correlation spectroscopy (Extended Data Figure 1c). LifA-Cy3B was concentrated to about 1 μM as described above. For determination of the labelling efficiency, the peak absorbance at A_568nm_ for Cy3B was measured and related to the protein peak absorbance at A_280nm_. The Cy3B labelling efficiency of LifA was 101 %.

### Vitrification of Samples and EM Data Acquisition

Gold grids (UltrAuFoil® R 0.6/1 300 mesh; *Quantifol*) were glow discharged in air at a pressure of 3.0×10^-1^ Torr at medium power for 120 s - 150 s with a Harrick Plasma Cleaner (PDC-002). The grids were plunge frozen in liquid ethane using a Vitrobot IV (*FEI*). The settings were 4 °C, 95 % humidity, blot time 5 s, blot force of 25, without drain and wait time. Movies were acquired with a Titan Krios G3 (*Thermo Fisher*) at 300 kV using EPU. For lymphostatin at pH 6.5, the Falcon III direct detector (*Thermo Fisher*) was used in linear mode at a nominal magnification of 75.000 and a total exposure of 73 e-/Å^2^ (Extended Data Table 1). At each stage position, movies were obtained from the central hole and the four closest surrounding holes using image shift without beam tilt compensation. Movies were stored in MRC-format and motion corrected and dose weighted with Motioncorr2 [35] during the data acquisition.

For lymphostatin at pH 4.0, movies were acquired at the same microscope with a Falcon IVi direct detector and a Selectris energy filter (*Thermo Fisher*) in counting mode with zero-loss imaging (slit width of 5 eV). Movies were obtained with the fast option of EPU that uses aberration free image shift (AFIS) [36] within a radius of 12 μm of the stage position. The holes were selected based on the plasmon image of a grid mesh [37] and the ice filter was adjusted for a narrow distribution of the ice thickness. The movies were acquired in EER format and pre-processed in a CryoSPARC Life session [38] including patch motion correction, dose weighting, patch CTF determination, blob picking, particle extraction and 2D-classification (Extended Data Table 1, Extended Data Figure 8).

### EM Image Processing and Model Building

The image processing and the initial ab-initio reconstructions were done with CryoSPARC v4.0 – 4.4 [38] using standard procedures. The particle images had intrinsic flexibility, preferential orientation and two conformations. Therefore, intense sorting with several rounds of 2D-classification, and 3D-classification with heterogeneous refinement or variability analysis were required to identify homogeneous subsets of the particles. Individual clusters and/or classes were refined with nonuniform refinement. With this strategy subsets of particles were identified, which were more isotropic in resolution than others. The Interpretability of the maps was judged by automated model building with ModelAngelo [39] based on the completeness of the models. The parameters of the image processing are summarized in Extended Data Table 1.

### Processing of lymphostatin at pH 4.0

Movies were processed with a CryoSPARC life session and particle locations were determined with the blob picker. Starting references were determined from a subset of 1.7 million extracted particle images. This subset was further divided into three subsets based on the appearance of 2D-class averages (small-, large- and medium sized particles). From each subset ab-initio reconstructions were obtained, and the data was further refined and sorted by heterogeneous refinement and 3D-variability analysis followed by nonuniform refinement. Eventually, maps were obtained for the full particle and both arms (GT-arm and delivery-arm). The three maps were aligned in respect to each other with Align-3D. In addition, the maps of the arms were mass-centred with ‘relion_image_handler’ and reimported to CryoSPARC. The 5 aligned maps served as references (Extended Data Figure 8a) for heterogeneous refinement of the full blob-picked set of particle images.

In total, 14,699,202 blob picked particle images were extracted during the life session. Three rounds of 2D-classification and class selection reduced the number of accepted particle images to 6,559,403. These particle images were heterogeneously refined with the previously determined starting references (Extended Data Figure 8a). 1,486,156 particle images grouped to class 1 and showed the full particle, while the other classes represented either the delivery arm or the GT-arm (Extended Data Figure 8b). Closer inspection of the classes suggested that delivery arm and GT-arm were parts of the full particle but were picked off-centre.

### Processing of Class 1

Particle images in class 1 were none-uniform refined, followed by 3D-variability analysis and 3D-Variability display into 5 clusters. All five clusters were none-uniform refinement (Extended Data Figure 8c). The resolution of the clusters ranged between 2.6 Å and 3.2 Å. The clusters were further evaluated by automated model building with ModelAngelo and orientation diagnostics. This identified cluster 5 as the best cluster with the most complete model (Extended Data Figure 8d).

### Processing of template picked particle images

To overcome the problem of off-centre picking, we re-determined particle locations by template picking. To generate a template, the 3D-map of cluster 5 was projected into 50 equally spaced directions. Template picking identified 13,373,262 particle locations, from which particle images were extracted.

The particle images were 2D-classified and the particles in the best classes were retained. This was followed by another round of 2D-classification and selection leaving 7,650,753 particle images in the data set. These particle images were classified by 3D-heterogeneous refinement using three different maps of the full particle as starting references. Most particles grouped to two references and represented conformation I (2,516,155 particle images) and conformation II (3,353,690 particle images). Subsequently, both conformations were processed separately.

### Processing of conformation I

The particle images in conformation I were heterogeneously refined into four classes. The best three classes were selected and again refined into three classes by heterogeneous refinement. This identified one well resolved class, and two with pathologies such as streaks or generally low resolution. The well-resolved class included 440,787 particles and was none-uniform refined using the uncropped boxes. At the end of the refinement the overall resolution was 2.7 Å (range by orientation diagnostic: 2.5-3.5 Å) with a sampling compensation factor of SCF=0.94 and a conical FSC Area Ratio is cFAR=0.30 (Extended Data Figure 8e).

Next, the particle images from the NU-refinement were restacked and imported into Relion 5.0 [40] using pyem [41] for converting between CryoSPARC “cs” and Relion “star”. A 3D-map was calculated from a subset of 10000 imported particle images with relion_reconstruct. This map was used for calculating a loose mask. Mask, map and imported particle images were the starting parameters for the subsequent 3D-classification without alignment into 3 classes. The largest and best resolved class contained 370,711 particles and was subjected to local refinement with relion_autorefine and blush regularization resulting in a map with 3 Å resolution.

For local refinement of the delivery domain, a loose mask of the tip of the arm was generated. The mask/region and the particle images from the previous consensus refinement were classified in 3D without alignment into 5 classes with a regularization parameter of T=10 and blush regularization followed by local refinement with relion_autorefine and blush regularization of the best class. The final map of the tip of the delivery domain contained 120,944 particles and had a resolution of 3 Å. Initial models of the consensus map, and the local refined maps were built with ModelAngelo from post-processed, and B-factor sharpened maps.

### Processing of Conformation II

The particle images in conformation II were subjected to another heterogeneous refinement into four classes in CryoSPARC. The best class included 1,637,694 particle images and was none-uniform refined using the uncropped particle images. The overall resolution at the end of the refinement was 2.4 Å (range by orientation diagnostic: 2.3-3.4 Å) with a sampling compensation factor of SCF=0.71 and a conical FSC Area Ratio of cFAR=0.19. As the cFAR was low, the data was subjected to another heterogeneous refinement into three classes, using the three copies of the map as starting references. All three classes were NU-refined. The largest class contained 812,767 particles and had an overall resolution of 2.5 Å (range by orientation diagnostic: 2.3-3.1 Å) with a sampling compensation factor of SCF=0.65 and a conical FSC Area Ratio of cFAR=0.33 (Extended Data Figure 8f). The other two classes were too anisotropic in resolution (range by orientation diagnostic 2.4 Å-9.1 Å, cFAR=0.02 and 2.5 Å -10.2 Å, cFAR=0.02) for further interpretation.

The particle images from the none-uniform refinement of the largest class were restacked in CryoSPARC and imported to Relion 5.0 [40] using pyem [41]. A 3D-map was calculated from a subset of 50,000 particle images of the imported stack with relion_reconstruct. This map was used for calculating a loose mask. Mask, map and imported particle stack were the starting parameters for subsequent 3D-classification without alignment into 3 classes. The largest and best resolved class contained 627,040 particle images and was locally refined with relion_autorefine and blush regularization resulting in a consensus map with 2.8 Å resolution.

For the local refinement of the delivery domain a spherical mask was generated *de novo*. The mask was centred at the tip of the delivery arm and had a diameter of 140 Å. This mask and the consensus refinement of conformation II were used as starting parameters for the focussed 3D-classification without alignment into 3 classes (T=8, blush regularization). The classes of the focussed classification showed somewhat different orientations of the delivery domain. The best resolved class was locally refined with relion_autorefine and blush regularization and had a resolution of 3.8 Å (182,154 particle images). Automated model building with ModelAngelo built 425 residues of the C-terminal part of the delivery domain beyond residues 2434. The ModelAngelo model together with the model of conformation I enabled modelling the rearrangement of the delivery domain by rigid body movement.

### Model building

A composite map of conformation I was generated using the map of the full particle from the blob picked particles (Extended Data Figure 8d) and the locally refined delivery domain of conformation I with the “Combine Focused Maps” tool in the program Phenix [42] where the focus switched from one map to the other as defined by residue 2200 in the model. The composite map of conformation II was generated similarly, with the relion consensus map of conformation II and the locally refined delivery domain of conformation II, switching at residue 2434.

Manual model building with Coot [43] was guided by automated model building via ModelAngelo [39] in the two maps generated above. Phenix [42] was used for real-space refinements. The final models are refined against the composited maps (Extended Data Table 2).

### Domain and linker assignment

Initial assignment of the GT-I domain, the protease domain and the DUF3491 followed the assignment in the INTERPRO database [16]. The GT-II domain, and the ART domain were identified with Foldseek [24]. The ribfind [44] plugin in ChimeraX [45] was used to identify the linkers as regions that did not cluster within a designated rigid body and connecting the designated domains.

### Fluorescence Lifetime Imaging Microscopy (FLIM)

HEK293T FlipIn cells (*ThermoFischer Scientific*) stably expressing the G protein-coupled receptor neurotensin receptor 1 linked to the fluorescent protein mNeonGreen, NTSR1-mNeonGreen [30, 46], were seeded into microfluidic chambers with 170 μm cover glass bottom (μ-Slide VI^0.5^, *IBIDI*) 3 days in advance. The cells grew at 37 °C to about 30 % confluency. To reduce background fluorescence, growth medium, DMEM (*ThermoFischer Scientific*) supplemented with 10% fetal bovine serum (FBS) (Bio&Sell), 1 % penicillin/streptomycin (*Sigma*) and, 3 μg/ml puromycin (*InvivoGen*), was exchanged for PBS buffer (DPBS, *ThermoFischer Scientific*) supplemented with 0.9 mM CaCl_2_ and 0.5 mM MgCl_2_ 1 h before the start of microscopic imaging. This starvation period stopped the expression of new NTSR1-mNeonGreen while matured GPCRs were transported to the cell membrane. 10 μl LifA-Cy3B (∼60 nM final concentration) was added in one entry port of the microfluidic chamber on the microscope to record the same cells before and after the addition of LifA-Cy3B. For overnight incubation LifA-Cy3B was added directly to the growth media in the microfluidic chamber. After 15 h incubation at 37 °C, the media was exchanged for supplemented PBS buffer. Cells were kept at 37 °C by a stage top incubator system (*OKO lab*) during the microscopic measurements.

Dual-colour fluorescence lifetime imaging microscopy was recorded with a confocal STED-FLIM microscope (ExpertLine, *Abberior Instruments*). Picosecond pulsed laser excitation was applied with 488 nm for mNeonGreen and 561 nm for Cy3B at 40 MHz repetition rate. A 60x water immersion objective with numerical aperture 1.2 (UPLSAPO 60XW, *Olympus*) allowed extended z-stack imaging. Fluorescence of mNeonGreen was detected in the spectral range from 500 to 550 nm. Fluorescence of Cy3B was detected in the combined spectral range from 580 to 630 nm and from 650 to 720 nm. Each spectral range was separated by a polarizing beam splitter cube to enable fluorescence anisotropy analysis. Photons were recorded by six single photon counting avalanche photodiodes with optimized time resolution (SPCM-AQRH-14-TR, *Excelitas*) using synchronized TCSPC electronics (SPC-154N, *Becker & Hickl*). Signals from different detectors could be combined if needed. Time binning was set to 21 ps to obtain high-resolution fluorescence lifetime decays.

### Fluorescence Lifetime Analysis

The software SPCImage NG (version 8.9, *Becker & Hickl*) was used for fitting the FLIM data. The at 561 nm and detected from 570 to 650 nm. Cells were kept at 37 °C during the measurements by a stage top incubator.

## Accession numbers for deposited maps

EMD-19982 (conformation I, consensus), EMD-19983 (conformation I, focused C-terminal part), EMD 19987 (conformation I, composite map), EMD-19984 (conformation II, consensus), EMD-19985 (conformation II, focused C-terminal part), EMD-19988 (conformation II, composite map). 9EUV and 9EUW are the PDB entries for the models of conformation I and II, respectively.

instrumental response function was determined from the data. Photon statistics were improved using a 2×2 pixel binning. Intensity thresholds were applied to exclude parts of the images with low background. Lifetime fitting was performed using maximum likelihood estimation for incomplete decay models, i.e., a monoexponential decay function for NTSR1-mNeonGreen or a biexponential decay function for LifA-Cy3B. For LifA-Cy3B, only the longer lifetime component was considered for FLIM-based cellular localization.

### Fluorescence Correlation Spectroscopy (FCS)

Fluorescence correlation spectroscopy was performed on the same STED-FLIM microscope (*Abberior*) using the 60x water immersion objective and pulsed excitation with 561 nm at 40 MHz. Data recording in FIFO mode was controlled by the software SPCM (*Becker & Hickl*). Autocorrelation functions were calculated and analysed by the software Burst_Analyzer (*Becker & Hickl*).

### Confocal Timelapse Imaging

Cellular uptake of LifA-Cy3B to HEK-293T cells in timelapse imaging over several hours was recorded with a different laser scanning microscope (LSM 980 Airy scan, *Carl Zeiss*) using continuous-wave lasers and an 60x water immersion objective. Fluorescence of mNeonGreen was excited with 488 nm and detected from 500 to 570 nm. Cy3B was excited

## Acknowledgements

Cryo-electron microscopy was carried out in the cryo-EM facility of the Julius-Maximilians-Universität Würzburg funded by the Deutsche Forschungsgemeinschaft (DFG, German Research Foundation – Projects INST 93/903-1 #359471283, INST 93/1042-1 #456578072, INST 93/1143-1 # 525040890). B.B. and M.G. acknowledge project funding by the DFG (Project Bo1150/18-1 #428774170). M.P.S. acknowledges strategic investment by the Biotechnology & Biological Sciences Research Council (BBS/E/RL/230002C). M.B. acknowledges funding by the State of Thuringia and the DFG for the confocal FLIM-STED microscope, Abberior Instruments (DFG project INST 1757/25-1 #411346541).

## Author contributions

M.G., T.R.; M.B., L.S. and B.B. developed the experimental design. M.G., R.S. and L.S conducted the experiments. C.K, T.R. and B.B. collected the data. B.B. processed the electron microscopic image data, L.S. und M.B. processed the light microscopic data. T.R. and M.G. built the molecular models. V.J.F., M.G. L.S. M.B. and B.B designed the figures. M.G., M.P.S., M.B. and B.B. interpreted the data. All authors contributed to writing and editing of the manuscript.

