## Extended Figures for "Lymphostatin: Structure of a large multi-functional virulence factor"

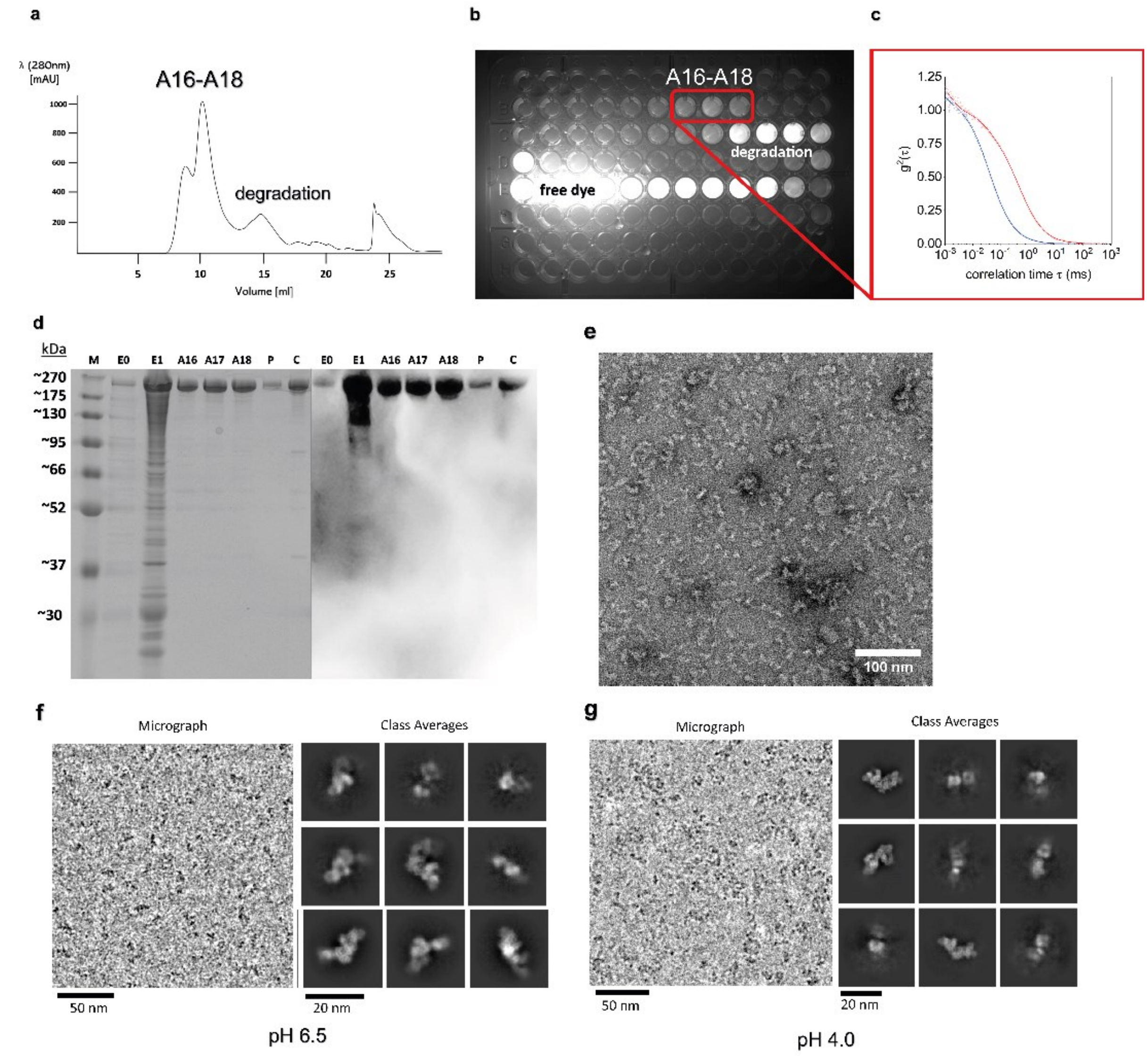

**Extended Data Figure 1 | Protein analytics after purification and concentration of full-length LifA. a,** Typical profile of size-exclusion chromatography of LifA applying main elution fraction of IMAC. Full-length LifA elutes after approx. 10 ml with three peak fractions (≈ 900 µl). At 15 ml a protein peak likely associated with degradation of full-length LifA is visible. **b,** After labelling LifA with fluorescent dye Cyanine3B maleimide (Cy3B), successful separation of free dye is visible on SEC elution plate under green excitation. **c,** Fluorescence correlation spectroscopy, FCS, of LifA-Cy3B (red curve) and Atto565-COOH (blue curve). TCSPC data were recorded with the STED-FLIM microscope (Abberior) using 561 nm pulsed laser excitation. Autocorrelation functions were calculated and analysed with the Burst_Analyzer software (Becker&Hickl). The diffusion time of ATTO 565-COOH was tD = 0.041±0.001 ms, the diffusion time of LifA-Cy3B was tD = 0.57±0.03 ms. Fitting the autocorrelation function of LifA-Cy3B indicated an additional small amount of unbound fluorophore in the range of less than 20 %. Based on an effective hydrodynamic radius of 0.7±0.07 nm for ATTO 565-COOH in aqueous buffer, the diffusion time of LifA-Cy3B corresponded to a mean diameter of about 19±2 nm for LifA, which is in good agreement with the size of a monomeric LifA protein in buffer. **d,** SDS-PAGE (left) and Western Blot (right) show purified LifA. M indicates marker with the highest marker band at 270 kDa (BlueEasy Prestained Protein Marker; Nippon Genetics). E0 and E1 indicate elution fractions from IMAC with E1 being used for subsequent SEC. A16, A17, A18 are peak fractions of full-length LifA Peak during SEC. P indicates pooled fractions A16, A17 and A18. C is the sample after concentration. Samples P and C were diluted for protein analytics to better compare samples before and after concentration. Full-length LifA migrates just under 270 kDa. **e,** Typical appearance of full-length LifA in negative stained samples (approx. 75 µg/ml). Particles show different shapes and sizes up to approx. 20 nm in line with an L-shaped particles of 365 kDa with different orientations. **f,** Lymphostatin at pH 6.5 –image data: Close-up of a motion corrected and dose weighted movie average. The average is low pass filtered to 1/(8 Å) for depiction. It represents approximately ¼ of the full movie area. The movie is taken with approximately 3.5 µm underfocus in linear mode. Particle images and selected class averages are shown at the same scale. **g,** Lymphostatin at pH 4.0 –image data: Close-up of a motion corrected and dose weighted movie average. The average is low pass filtered to 1/(8 Å) for depiction. The close-up represents approximately ¼ of the full movie area. The movie is taken with approximately 2.0 µm underfocus as zero-loss image in counting mode. Particle images and selected class averages are shown at the same scale.

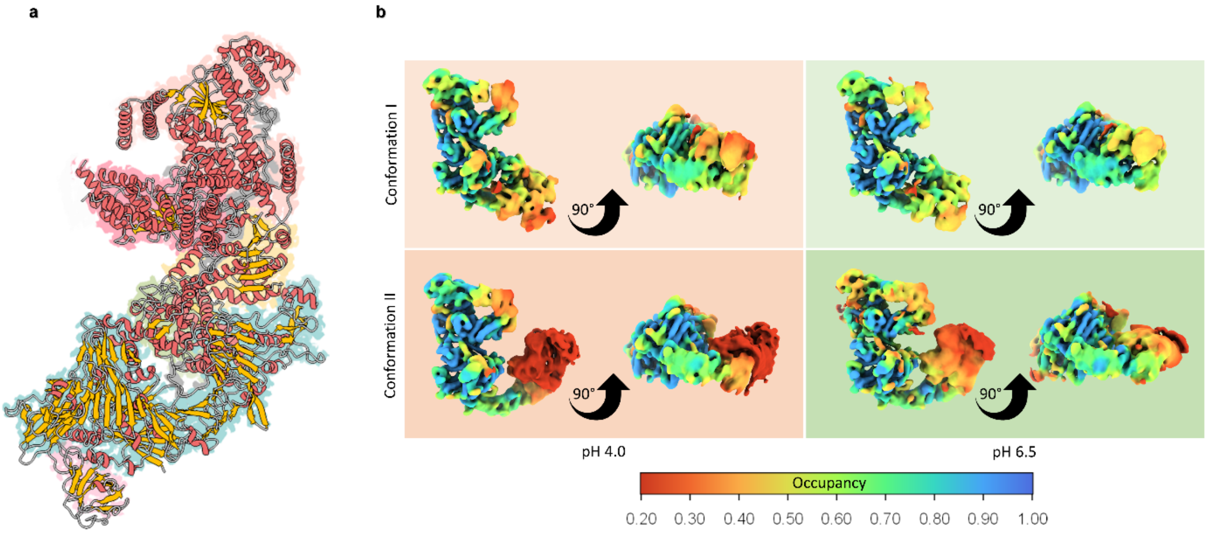

**Extended Data Figure 2 | LifA shows two structural distinctive arms and two coexisting conformations**. a, The surface of LifA is shown with domains depicted according to the used colour code. Overlayed, α-helices are shown in red, β-sheets in orange, and loops in white. The N-terminal arm shows an α-helix rich secondary structure that characterize the GT-I (peach), GT-II (red) and protease domain (yellow). The C-terminal arm shows an extended stretch of β-sheets observable throughout the delivery domain (blue) as well as the ART domain (pink). b, Conformation I and II coexist at pH 4.0 and 6.5: Surface representations of conformation I (upper panels) and conformation II (lower panels) scaled and coloured with occupy. Both conformations were observed at pH 4.0 (red background) and pH 6.5 (green background) in independent data sets. All maps are aligned in respect to each other. The map of conformation I at pH 4.0 was low pass filtered to 8 Å (relion_image_handler) and used as reference map for amplitude scaling of the other maps (relion_image_handler). The grey values of the resulting maps were adjusted with OccuPy (Extended Data Figure References 1) and the surface of the maps are coloured with the relative occupancy. In this context, low occupancy suggests more structural variability of a region. All maps are shown with the same threshold and were scaled with the same parameters in Occupy. All maps are shown in two perpendicular views.

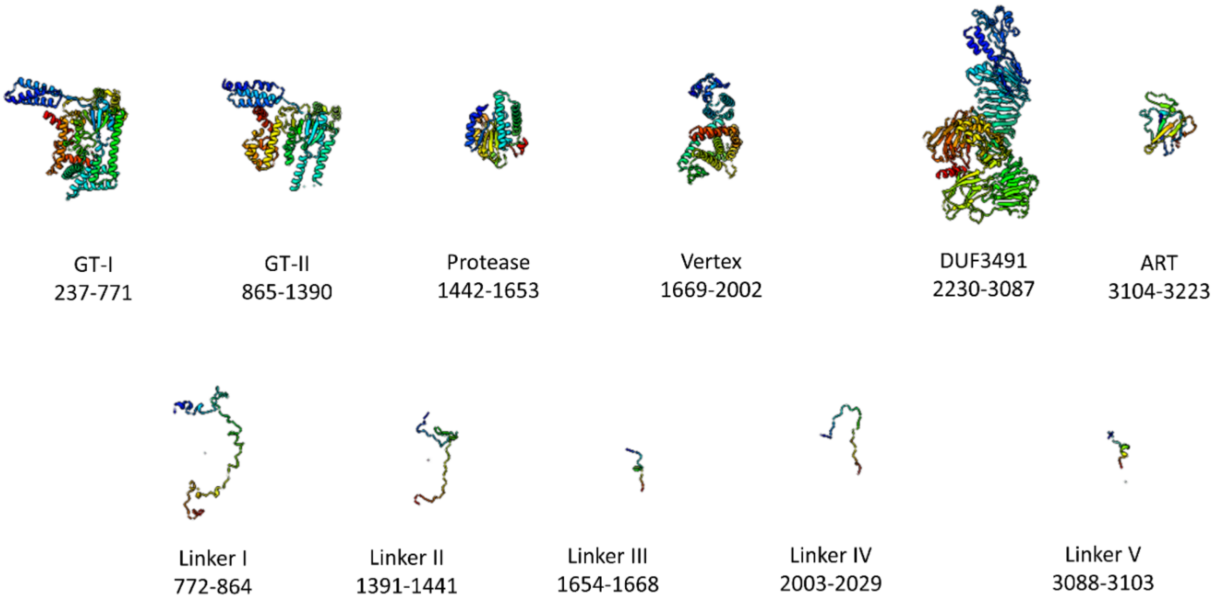

**Extended Data Figure 3 | Domains and linkers in lymphostatin**. The linkers and domains are coloured in rainbow from blue (N-terminus of domain/linker) to red (C-terminus of domain/linker). The orientation in which the domain is shown is arbitrary and places the N-terminus at the top left. All domains and linkers are shown at the same scale.

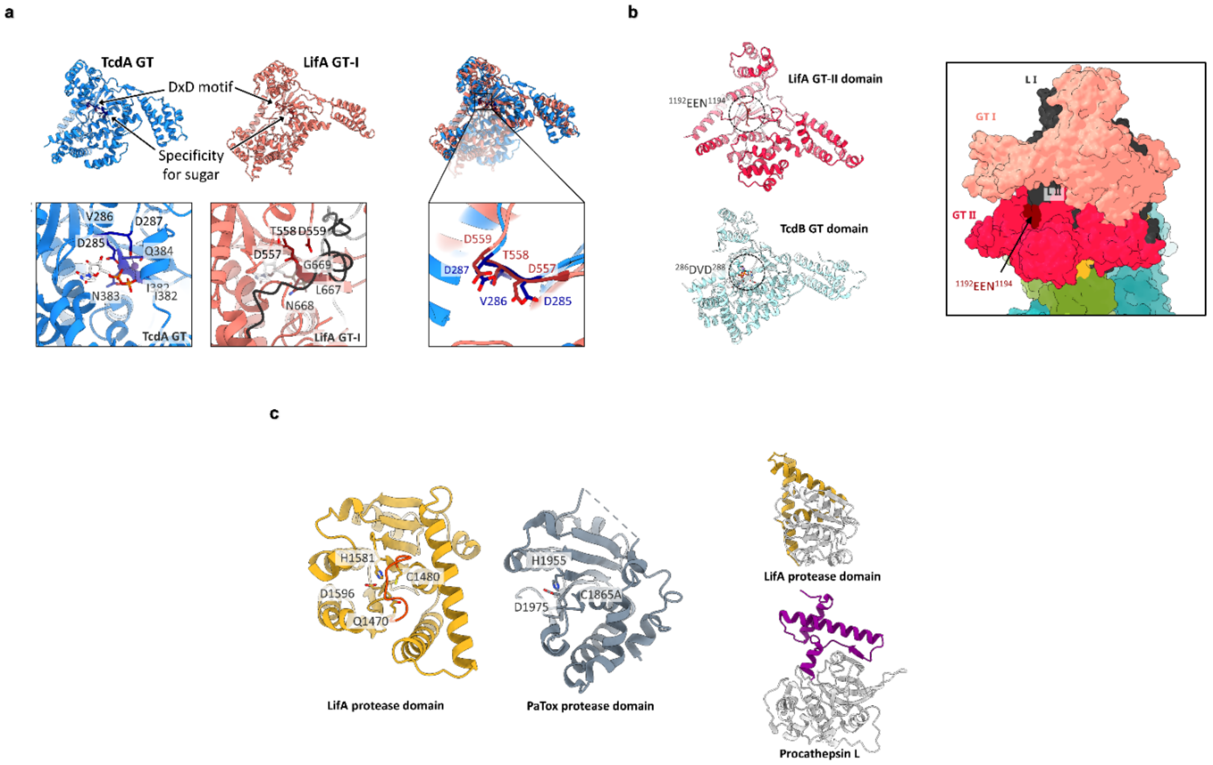

**Extended Data Figure 4 | Detail views of N-terminal domains of lymphostatin. a,** LifA GT-I domain is structurally related to the clostridial toxin TcdA GT-domain. LifA GT-I domain (*peach*) exhibits both a DXD motif as well as a specificity for sugar with a similar architecture to LCTs TcdA GT domain (blue).

Overlay of LifA GT-I domain (*peach*) and TcdA GT-domain (*blue*). The characteristic DXD motif (residues 557-559) in the active site of GT-I is very conserved between LifA (*peach*) and TcdA (*blue*). In contrast to TcdA/B (which show a valine residue), LifA GT-I has a threonine residue edged by two aspartic acids. Comparison between crystal structure (PDB ID: 4DMW, Malito et al., 2012) of the GT domain (*blue*) of *Clostridium difficile* toxin A (TcdA) in complex with UDP and Manganese (*left*). On the right, the UDP of the original TcdA-GT structure is superposed with the equivalent presumptive sugar binding region of the LifA GT-I domain (right, *peach*). Here, the active site is blocked by linker I (*black*) formed by residues 532 – 542 which fold back into the putative position of the sugar substrate. **b,** LifA GT-II domain shows an exposed EEN motif. Comparison of LifA GT-II domain (*red*) to *Clostridioides difficile* toxin B (TcdB) crystal structure (only one chain is shown) in complex with UDP and neomycin (PDB ID: 7LOV, Harijan et al., 2021, *light blue*), which was identified as the closest structural homologue by Foldseek Search Server. The DxD motif of TcdB is replaced with an EEN motif in LifA GT-II. The surface of LifA is shown focused on GT-I (*peach*) and GT-II (*red*). The EEN motif (residues 1192-1194, *brown*) of the LifA GT-II domain is exposed and fully accessible. **c,** The active site of the LifA protease domain is not accessible. The comparison of LifA protease domain (left, *yellow*) to the protease domain of PaTox (PDB ID: 6HV6, Bogdanovich et al., 2019, right*, blue grey*) from *Photorhabdus asymbiotica* shows high degree of conservation to other C58-proteases. The active site of the LifA protease domain consists of the catalytic triad (C1480, H1581 and D1596). Additionally, Q1470 is functionally relevant for stabilizing reaction intermediates. A loop (*orange*) connecting functional residues Q1470 and C1480 is blocking the entry to the active site of LifA protease domain. LifA (top) shows long crossing helices (*yellow*) in its protease domain reminiscent of pro-domains of papain-like proteases like human procathepsin L (PDB ID: 1CS8, Cygler and Coulombe, 1999, bottom, long crossing helices depicted in purple) that regulate protease activity.

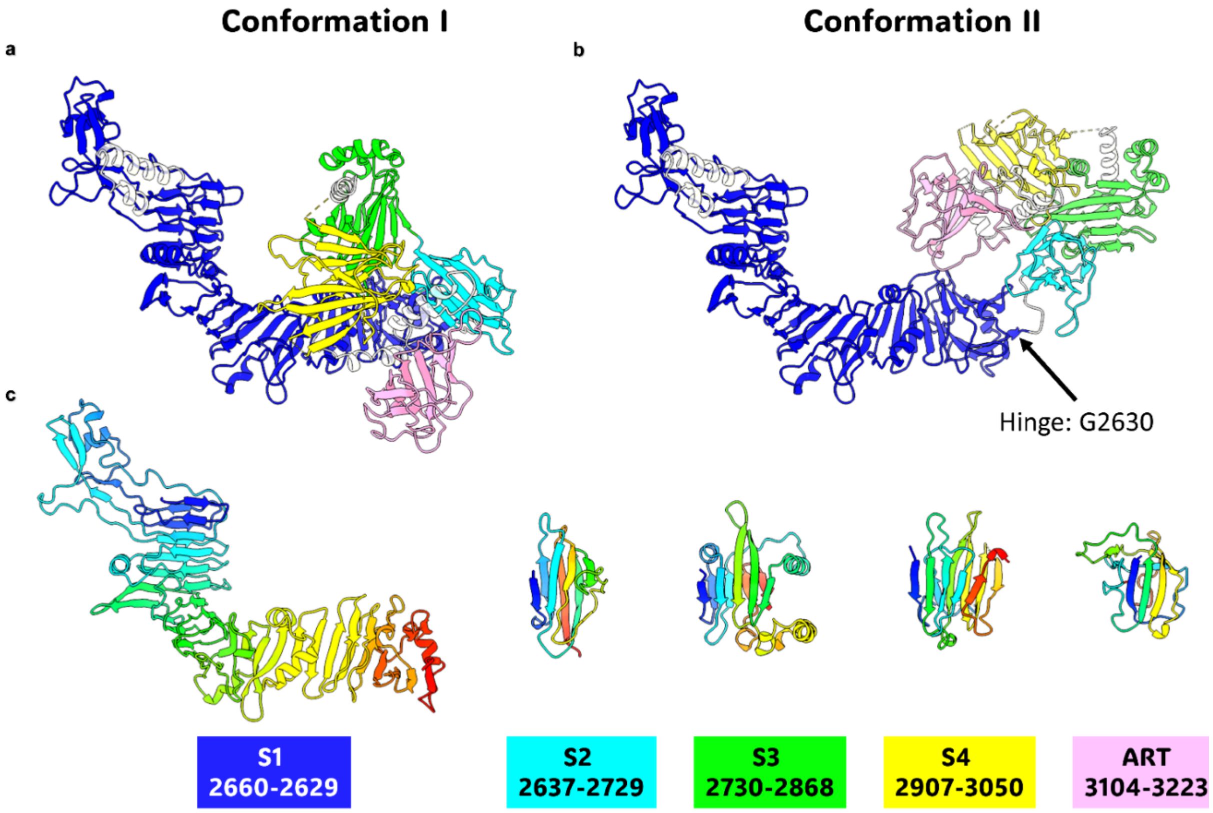

**Extended Data Figure 5 | The delivery domain has four subdomains and is differently arranged in conformation I and II**: **a,** and **b,** Models of the C-terminal arm (2660-3232) in conformation I and conformation II. The delivery domain has four ß-sandwich subdomains S1-S4 which are coloured blue (S1), cyan (S2), green (S3) and yellow (S4) and is followed by the ART domain (*pink*). Conformation I and Conformation II differ by a rigid body rotation of S2-S4 and the ART domain around G2630 (indicated by an arrow) relative to S1. **c,** shows the subdomains S1-S4 and the ART domain in arbitrary orientations placing the N-terminus close to the top left. The domains are coloured in rainbow from their N-terminus (*blue*) to their C-terminus (*red*). The names of the (sub)-domains together with their residues are given below. The background of the label is coloured the same as the subdomain in a and b. All models are shown at the same scale.

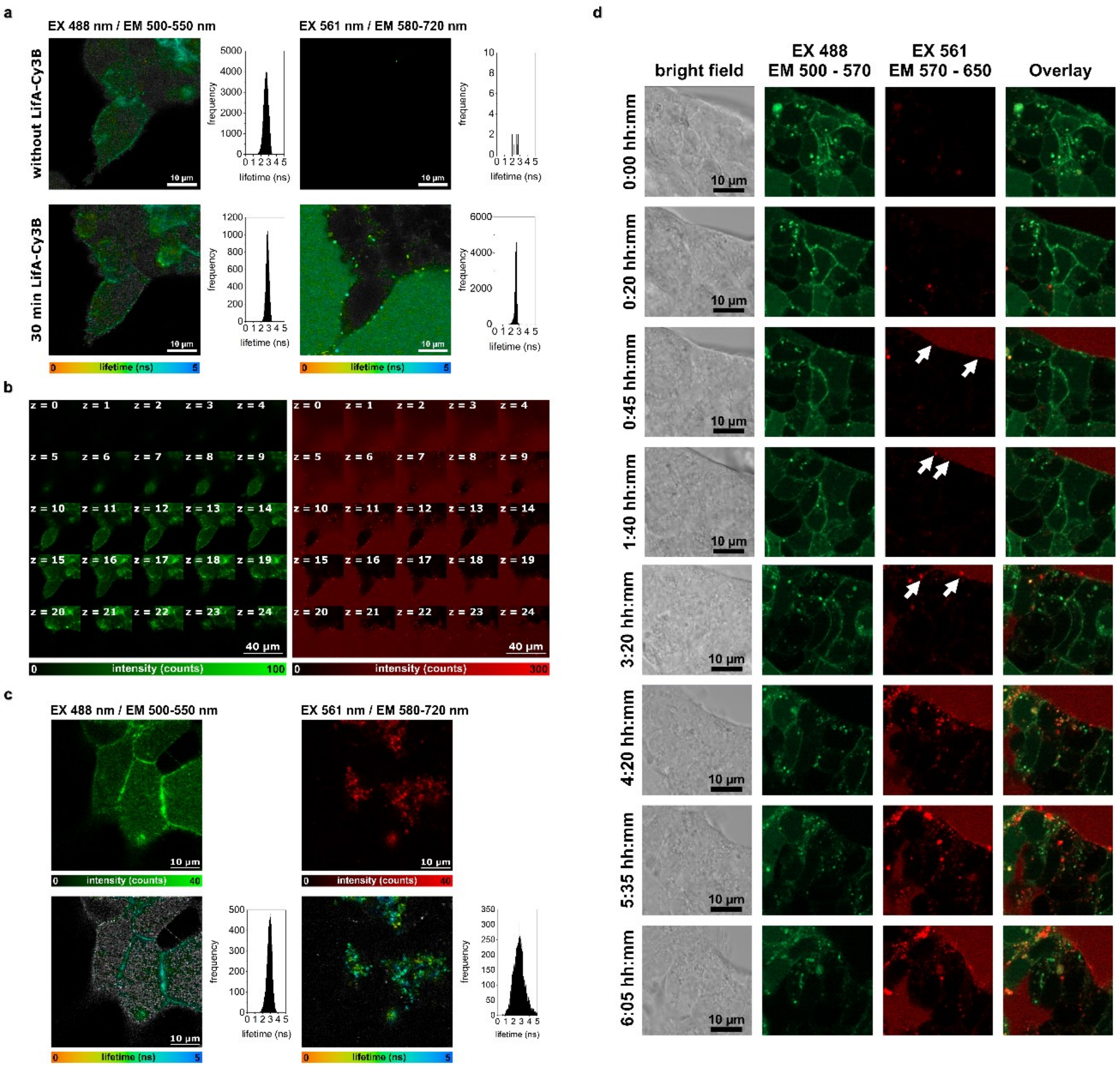

**Extended Data Figure 6 | Fluorescence microscopy of Cy3B-labelled LifA. a,** Spectrally separated FLIM images of HEK-293T cells (related to composite Figure 4a in the manuscript), before (upper panels) and 30 min after addition of LifA-Cy3B (lower panels). Left panels show NTSR1-mNeonGreen with laser excitation at 488 nm and detection between 500 and 550 nm, right panels show LifA-Cy3B with laser excitation at 561 nm and detection between 580 and 720 nm. Note that photons in the spectral range between 630 and 650 nm are not detected due to optical filters. Fluorescence lifetime distributions of intensity-selected pixels are shown in the adjacent histograms on the right side of each image. For LifA-Cy3B, a biexponential decay fit was required comprising a short lifetime of less than 600 ps assigned to autofluorescence and a longer lifetime around 2.7 ns from Cy3B. The long lifetime component is shown here in the FLIM images and histograms. FLIM images were recorded with the STED-FLIM microscope (Abberior) and analysed with the software SPCImage (*Becker&Hickl*). **b,** Complete series of z-stack FLIM images related to Figure 4A in the manuscript. The z-stack war measured 30 min after addition of LifA-Cy3B to the HEK-293T cells, without buffer exchange. In total, 25 different z layers with a distance of 1 mm were recorded. The images at z=14 mm were chosen for Figure 4A. Left panels show NTSR1-mNeonGreen with laser excitation at 488 nm and detection between 500 and 550 nm, right panels show LifA-Cy3B with laser excitation at 561 nm and detection between 580 and 720 nm. Intensities are given as counts per pixels, without background subtraction. Images were recorded with the STED-FLIM microscope (*Abberior*). **c,** Spectrally separated intensity (upper panels) and FLIM images (lower panels) of HEK-293T cells related to Figure 4C in the manuscript, after 15 h incubation with LifA-Cy3B in growth medium. LifA-Cy3 containing solution was replaced by PBS buffer 1 h before the recording. Left panels show NTSR1-mNeonGreen with pulsed laser excitation at 488 nm and detection between 500 and 550 nm, right panels show LifA-Cy3B with pulsed laser excitation at 561 nm and detection between 580 and 720 nm. Note that the spectral range between 630 and 650 nm is not detected due to optical filters. Fluorescence lifetime distributions of intensity-selected, false-coloured pixels are shown in the adjacent histograms on the right side. For LifA-Cy3B, a biexponential decay fit was required comprising a short lifetime of less than 600 ps assigned to autofluorescence and a longer lifetime around 2.7 ns from Cy3B. This long lifetime component is shown here in the FLIM images and histograms. Images were recorded with the STED-FLIM microscope (*Abberior*). **d,** Dual-colour time lapse imaging of HEK-293T cells with NTSR1-mNeonGreen (green channel) following addition of LifA-Cy3B (red channel). Grey images on the left side are the related transmission images. The overlay of NTSR1-mNeonGreen and LifA-Cy3B are shown on the right side. The time series was recorded with the LSM 980 microscope (*Zeiss*) using in slightly different spectral ranges for fluorescence detection. White arrows highlight the occurrence of bright LifA-Cy3B clusters on the plasma membranes, starting about 45 min after addition of LifA-Cy3B. These bright LifA-Cy3B clusters increased in size and brightness over time on the membrane, but also started to appear within the cells after about 4 hours. Cells changed their shape over time and the focal plane might have shifted, so that the later images include larger areas of cell medium with freely diffusing LifA-Cy3B outside the cells.

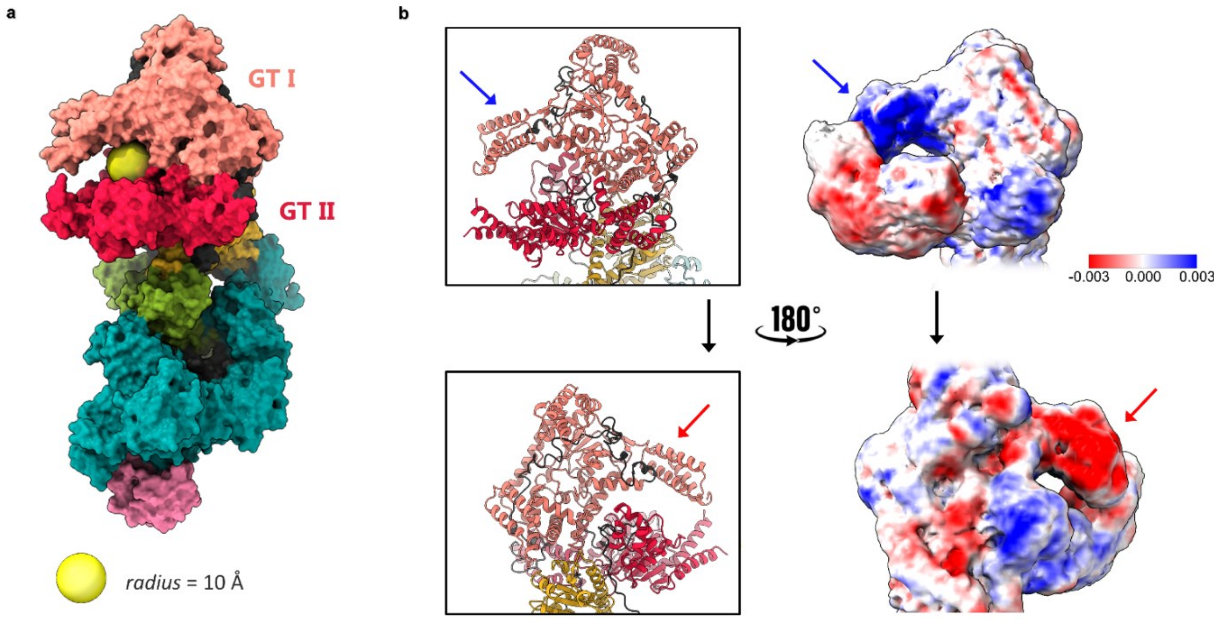

**Extended Data Figure 7 | The binding site of GT-II is enclosed by mobile helices from GT-I. a,** Surface representation coloured by the domains of LifA. The arrangement of the N-terminal helices of GT-I (*peach*) and a helix bundle of GT-II (*red*) give rise to a spherical cavity with a radius of approximately 10 Å (*yellow sphere*) for possible pre-binding of potential substrates. **b,** The models of GT-I and GT-II are depicted on the left, and a variability analysis (right) shows flexibility of N-terminal helices from functional GT-I domain. The corresponding helix regions are marked by a blue, or red, respectively, arrow**.**

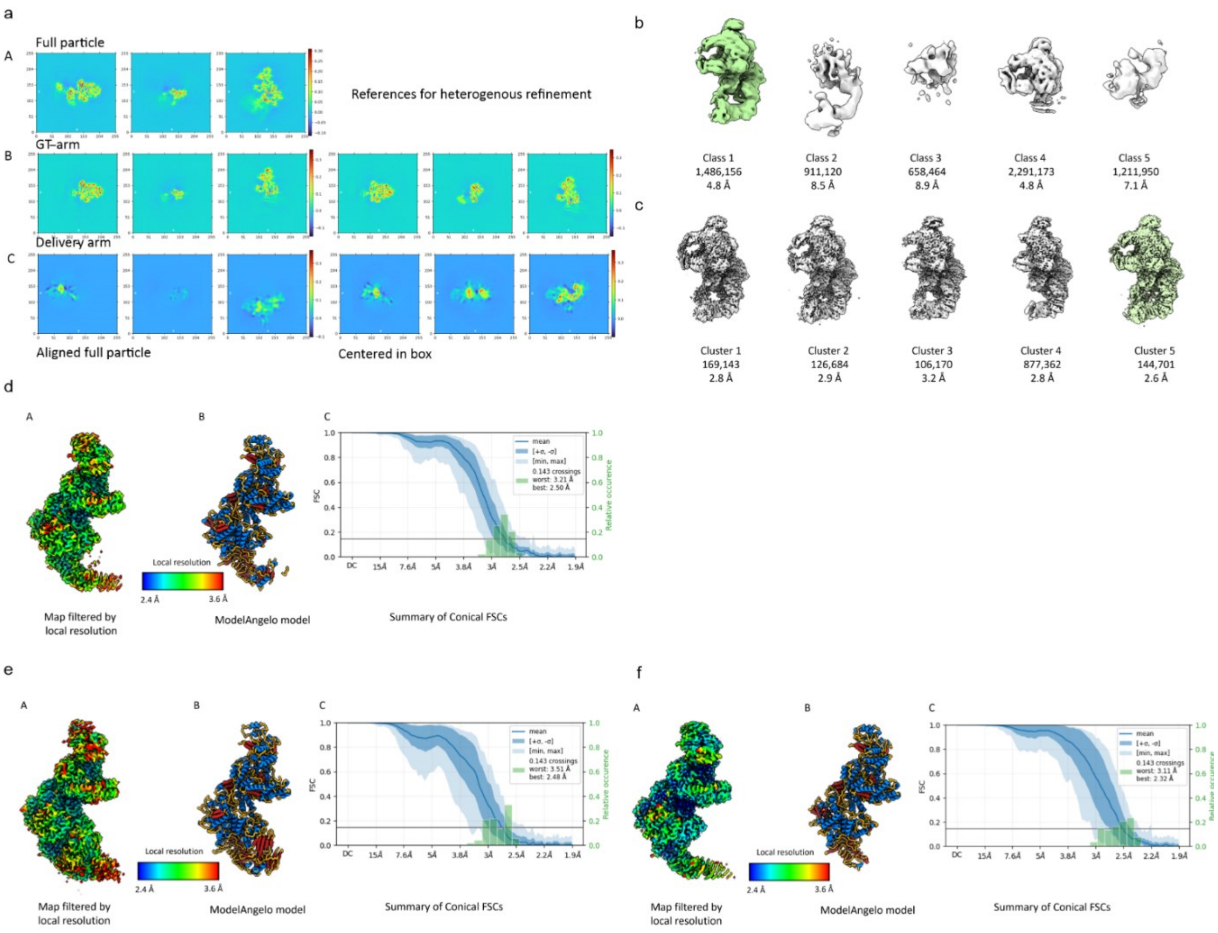

**Extended Data Figure 8 | Image processing. a,** Perpendicular slices through the starting references derived from different subsets of particle images: The starting references comprised A) the full particle, B) the GT-arm and C) the delivery arm. The maps of the arms were rotationally and translationally aligned to the full particle (left) or additionally mass-centred (right). **b,** Heterogenous refinement of the full data set - Surface representations of the final classes. For each class, the number of particle images is quoted and the nominal resolution. The heterogeneous refinement was done with Fourier cropped images. The Nyquist frequency is at 1/4.8 $Å^{-1}$. The image processing was continued with the particle images of class 1 (*green*). **c,** 3D-variability analysis of class 1 followed by NU-refinement of each cluster– Surface representation of the none-uniform refined clusters: The 3D-variability analysis was done with 3 components followed by 3D Variability display in cluster mode. Each cluster was none uniform refined. The surface representations of the final maps are not sharpened. For each cluster the number of particle images and the nominal resolution is quoted below. The best resolved cluster is cluster 5 (*green*). This is judged by the nominal resolution, the completeness of automated model building with ModelAngelo and the output of orientation diagnostics. The 3D-map of cluster 5 was used to generate templates for template picking**.**

**d-f)** Validations of maps for LifA.: d) derived from cluster 5; e) derived from conformation I of template picked particles; f) derived from conformation II of template picked particles Each validation shows A) The surface representation of the map filtered by local resolution and B-factor sharpened with B=-50 $Å^2$. The surface is coloured according to the local resolution. As indicated by the colour key. B) The model built automatically by ModelAngelo. The strands are shown in red and the helices in blue. C) The summary of the conical FSCs determined with the 3D Orientation diagnostic. In **d)** the anisotropic resolution ranges from 2.5-3.2 Å with a conical FSC Area Ratio of cFAR=0.42 and a sampling compensation Factor of SCF=0.80. In **e)** the anisotropic resolution ranges from 2.5-3.5 Å with a conical FSC Area Ratio of cFAR=0.30 and a sampling compensation Factor is SCF=0.94. In **f)** the anisotropic resolution ranges from 2.3-3.1 Å with a conical FSC Area Ratio of cFAR=0.33 and a sampling compensation Factor of SCF=0.65.

**Extended Data Table 1 | Data collection parameters for LifA and image processing of consensus maps**

| Sample | Lymphostatin pH 4.0 | | Lymphostatin pH 6.5 | |
|---|---|---|---|---|
| Electron microscope | Krios G3 with X-FEG (Thermo Fisher), Cs=2.7 mm, HT=300 kV | | | |
| Camera | Falcon IVi | | Falcon III | |
| Camera mode | counting | | linear | |
| Energy filter | Selectris (5eV slit) | | none | |
| Movie format | EER | | MRC, 40 fractions | |
| C2/Spot Size | 70 µm / Spot 5, nanoprobe | | | |
| Objective Aperture | none | | 100 µm | |
| Magnification | 130,000 (EFTEM) | | 75,000 | |
| Calibrated Pixel Size | 0.945 Å/Px | | 1.064 Å/Px | |
| Beam diameter | 1.3 µm | | 1 µm | |
| Total Exposure | 70 $e^{-}$/Å² | | 73 $e^{-}$/Å² | |
| Exposure time | 6.1 s | | 5.2 s | |
| Exposures per hole | 1 | | 1 | |
| Exposures per stage position | Unknown<br>Ca. 60-150 | | 5 | |
| Acquisition at stage position | Fast (AFIS) | | Image shift without beam tilt compensation | |
| Target range of under focus | 0.5-1.4 µm | | 1.4-2.6 µm | |
| Movies | 41392 | | 19325 | |
| Motion correction and Dose weighting | CryoSparc 4.4 Life patch-motion | | Motion Cor2 | |
| Particle Images at start | 7,650,753 | | 1,206,880 | |
| Particle images in consensus map | Conf.I<br>440,787 | Conf. II<br>812,767 | Conf. I<br>410,717 | Conf. II<br>407,590 |
| Overall resolution | 2.7 Å | 2.5 Å | 3.4 Å | 3.4 Å |
| Resolution range | 2.5-3.5 Å | 2.3-3.1 Å | 3.2-4.0 Å | 3.1-7.9 Å |
| Sampling Compensation Factor | 0.94 | 0.62 | 0.86 | 0.66 |

**Extended Data Table 2 | Cryo-EM data collection, refinement and validation statistics**

| | LifA conf.1 (EMDB-19987) (PDB 9EUV) | LifA conf.2 (EMDB-19988) (PDB 9EUW) |
|---|---|---|
| **Data collection and processing** | | |
| Magnification | 130,000 | 130,000 |
| Voltage (kV) | 300 | 300 |
| Electron exposure (e–/Å$^2$) | 70 | 70 |
| Defocus range (μm) | 0.5-1.4 | 0.5-1.4 |
| Pixel size (Å) | 0.946 | 0.946 |
| Symmetry imposed | - | - |
| Initial particle images (no.) | 13,373,262 | 13,373,262 |
| Final particle images (no.) | 440,787 | 627,040 |
| Map resolution (Å) | 2.7 | 2.8 |
| FSC threshold | 0.143 | 0.143 |
| Map resolution range (Å) | 2.5-3.5 | 2.3-3.1 |
| **Refinement** | | |
| Initial model used (PDB code) | - | - |
| Model resolution (Å) | 2.9 | 2.8 |
| FSC threshold | 0.5 | 0.5 |
| Model resolution range (Å) | | |
| Map sharpening *B* factor (Å$^2$) | | |
| Model composition | | |
| Non-hydrogen atoms | 22287 | 22324 |
| Protein residues | 2796 | 2801 |
| Ligands | - | - |
| *B* factors (Å$^2$) | | |
| Protein | 105 | 106 |
| Ligand | - | - |
| R.m.s. deviations | | |
| Bond lengths (Å) | 0.002 | 0.003 |
| Bond angles (°) | 0.438 | 0.545 |
| Validation | | |
| MolProbity score | 1.57 | 1.92 |
| Clashscore | 5.99 | 8.22 |
| Poor rotamers (%) | 1.73 | 2.61 |
| Ramachandran plot | | |
| Favored (%) | 97.7 | 97.1 |
| Allowed (%) | 2.3 | 2.8 |
| Disallowed (%) | 0.0 | 0.0 |

**Extended Data References**

Forsberg BO, Shah PNM, Burt A. A robust normalized local filter to estimate compositional heterogeneity directly from cryo-EM maps. Nat Commun. 2023;14(1):5802. Published 2023 Sep 19. doi:10.1038/s41467-023-41478-1

N, Malito E, Biancucci M, et al. The structure of Clostridium difficile toxin A glucosyltransferase domain bound to Mn2+ and UDP provides insights into glucosyltransferase activity and product release. FEBS J. 2012;279(17):3085-3097. doi:10.1111/j.1742-4658.2012.08688.x

Paparella AS, Aboulache BL, Harijan RK, Potts KS, Tyler PC, Schramm VL. Inhibition of Clostridium difficile TcdA and TcdB toxins with transition state analogues. Nat Commun. 2021;12(1):6285. Published 2021 Nov 1. doi:10.1038/s41467-021-26580-6

Bogdanovic X, Schneider S, Levanova N, et al. A cysteine protease-like domain enhances the cytotoxic effects of the Photorhabdus asymbiotica toxin PaTox. J Biol Chem. 2019;294(3):1035-1044. doi:10.1074/jbc.RA118.005043

LaLonde JM, Zhao B, Janson CA, et al. The crystal structure of human procathepsin K. Biochemistry. 1999;38(3):862-869. doi:10.1021/bi9822271
